# WASTER: Practical *de novo* phylogenomics from low-coverage short reads

**DOI:** 10.1101/2025.01.20.633983

**Authors:** Chao Zhang, Rasmus Nielsen

## Abstract

The advent of affordable whole-genome sequencing has spurred numerous large-scale projects aimed at inferring the tree of life, yet achieving a complete species-level phylogeny remains a distant goal due to significant costs and computational demands. Traditional species tree inference methods, though effective, are hampered by the need for high-coverage sequencing, high-quality genomic alignments, and extensive computational resources. To address these challenges, this study introduces WASTER, a novel *de novo* tool for inferring shallow phylogenies directly from short-read sequences. WASTER employs a k-mer based approach for identifying variable sites, circumventing the need for genome assembly and alignment. Using simulations, we demonstrate that WASTER achieves accuracy comparable to that of traditional alignment-based methods, even for low sequencing depth, and has substantially higher accuracy than other alignment-free methods. We validate WASTER’s efficacy on real data, where it accurately reconstructs phylogenies of eukaryotic species with as low depth as 1.5X. WASTER provides a fast and efficient solution for phylogeny estimation in cases where genome assembly and/or alignment may bias analyses or is challenging, for example due to low sequencing depth. It also provides a method for generating guide trees for tree-based alignment algorithms. WASTER’s ability to accurately estimate shallow phylogenies from low-coverage sequencing data without relying on assembly and alignment will lead to substantially reduced sequencing and computational costs in phylogenomic projects.

## 1 Introduction

Whole-genome sequencing has recently replaced single marker sequencing as the dominant method for estimating phylogenies, and has been the focus of many recent large-scale efforts for a broad set of species [1–10]. Nevertheless, these efforts are still short of realizing complete species-level phylogenies for most clades. Major obstacles include the costs of high-coverage sequencing, extensive human effort in running species tree inference pipelines, and substantial computational resource demanded [11]. In addition, alignment challenges may possibly affect the results in downstream phylogenetic analyses [12].

Traditional species tree inference pipelines typically involve several key steps: assembling reads into genomes, aligning orthologous genomic sequences, optionally reconstructing gene trees from aligned genomes, and inferring the species tree from gene trees or directly from the aligned genomes (Fig. 1a). The recently developed CASTER improves the efficiency of species tree inference in the presence of incomplete lineage sorting (ILS) by enabling direct inference from aligned genomes [13]. While CASTER helps alleviate the computational bottleneck from alignment to species tree, it still depends on genome assembly and genome alignment, which become new bottlenecks. Additionally, the necessity for high coverage sequencing persists, with traditional pipelines usually requiring coverage above 30X for reliable genome assemblies, imposing financial challenges [14, 15]. Additionally, Progressive Cactus, the leading tool for aligning a large number of genomes, relies on a user-specified guide-tree [16]. The use of a guide-tree for alignment may potentially lead to circular inferences when subsequently estimating the tree using the very same alignment. These factors underscore the need for accurate alignment-free and assembly-free species tree inference methods.

**Fig. 1.**
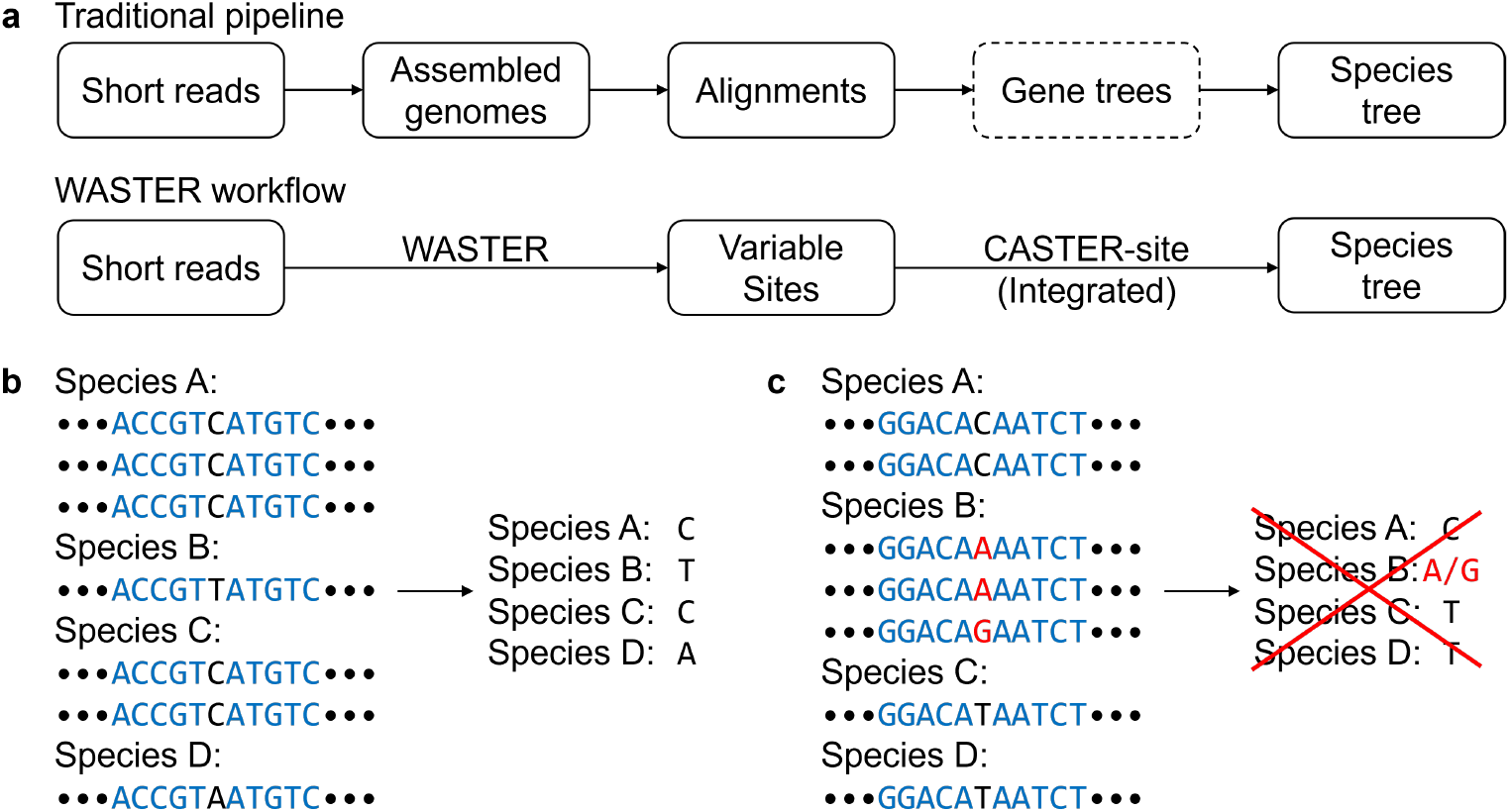
WASTER workflow. a, Overview of the WASTER workflow compared to the traditional species tree inference pipeline. **b**, WASTER indentifies variable sites by detecting k-mers that differ only at the central base. **c**, If two variants of a k-mer are found within the same species, WASTER removes the entire aligned site.

Several assembly-free methods have been proposed [17–22], but few are suitable for inferring trees from reads covering entire eukaryotic nuclear genomes. Some methods rely on mapping reads to reference sequences, thus requiring alignments of annotated orthologous sequences as input [21]. This limits their applicability, especially in the absence of closely related reference genomes. Other methods construct a “pangenome graph” encompassing all taxa, but this strategy does not scale well to genome-scale data [22]. Alternatively, several k-mer-based methods compute pairwise evolutionary distances and subsequently invoke distance-based algorithms to infer species trees [17– 20]. However, these methods struggle to infer accurate phylogenies for challenging trees with short internal branches.

In this article, we introduce WASTER (Without Alignment/Assembly Species Tree EstimatoR), a novel *de novo* tool that infers shallow phylogenies directly from short reads. WASTER operates by initially calling variable sites using k-mers from raw sequencing reads and then employs CASTER-site to infer the species tree from the matrix of variable sites (Fig. 1a). We evaluate the method extensively using simulations and applications to real data and show that it substantially outperforms other alignment-free methods and is as accurate as state-of-the-art alignment-based methods, even when assuming that the genome assembly and alignment are error free.

## 2 Results

### 2.1 WASTER: k-mer-based variable site identification

The variable site identification mechanism within WASTER involves a k-mer based approach. Initially, WASTER decomposes input reads or genomes into k-mers of a specified length *k*, where *k* is an odd number (19 by default). The process then groups these k-mers by matching all positions except the central one. For a given group of k-mers, if the k-mers from each species are identical, including the central position, then the central position of this k-mer group is designated as one variable site (Fig. 1b). In contrast, if any two k-mers from the same species differ, the entire group is disregarded, reducing the possibility of paralogy, alignment errors, or sequencing errors. This approach, however, results in the exclusion of heterozygous sites (Fig. 1c). Following the variable site identification, the concatenated variable sites are utilized as input for CASTER-site, which undertakes the inference of the species tree. Due to WASTER’s inability to distinguish between the forward and reverse strands, it employs the T92 substitution model [23], a simplified model that does not account for strand bias.

### 2.2 Evaluation in simulations

#### 2.2.1 SR201 simulated alignments

We first evaluated the performance of WASTER versus alignment-based methods using the SR201 dataset which comprises simulated alignments from 201 species with the heights of species trees similar to that observed in intrafamilial or intraordinal trees in birds and mammals [13]. This dataset incorporates many of the intricacies of molecular evolution. To avoid unrealistic strict molecular clocks, this dataset allows variation in generation time and population size across different branches of the species tree. Simulations were conducted under the Hudson model [24] for recombination and the GTR substitution model [25], with parameters adjusted for varying mutation rates across the genome. Notably, the equilibrium nucleotide frequencies differ from the no strand-bias model assumed by WASTER. We juxtaposed WASTER with several leading alignment-based methods:

1. RAxML-ng [26], a maximum-likelihood approach, widely used for reconstructing concatenated sequence tree yet statistically inconsistent [27],
2. SVDQuartets [28], an invariant-based method maintaining statistical consistency under ILS,
3. CASTER-site [13].

Although providing true simulated alignments as inputs, rather than assembled genomes from simulated reads, disproportionately favored alignment-based methods, WASTER still demonstrated accuracy on par with them. Across all experiments, WASTER’s species tree branch estimation error as measured by bipartition false negatives (FN) averaged at 3.9 branches, significantly lower (*p <* 10^*−*5^) than RAxML-ng (5.1 branches) and SVDQuartets (4.7 branches), but higher (*p <* 10^*−*15^) than CASTER-site (2.6 branches). Moreover, WASTER’s total runtime was comparable to the time required for species tree estimation from alignments alone in traditional workflows, being 5.1 times faster than RAxML-ng and 3.7 times faster than SVDQuartets (Fig. 2a). WASTER uses the integrated CASTER-site to infer species trees directly from sequence data. To evaluate the benefit of using CASTER-site over other methods in this context, we also tested an alternative approach in which variable sites extracted by WASTER were fed into SVDQuartets for species tree inference. Results show that default WASTER with CASTER-site yields higher accuracy than the combination of WASTER with SVDQuartets, highlighting the advantage of CASTER-site (Fig. A1).

**Fig. 2.**
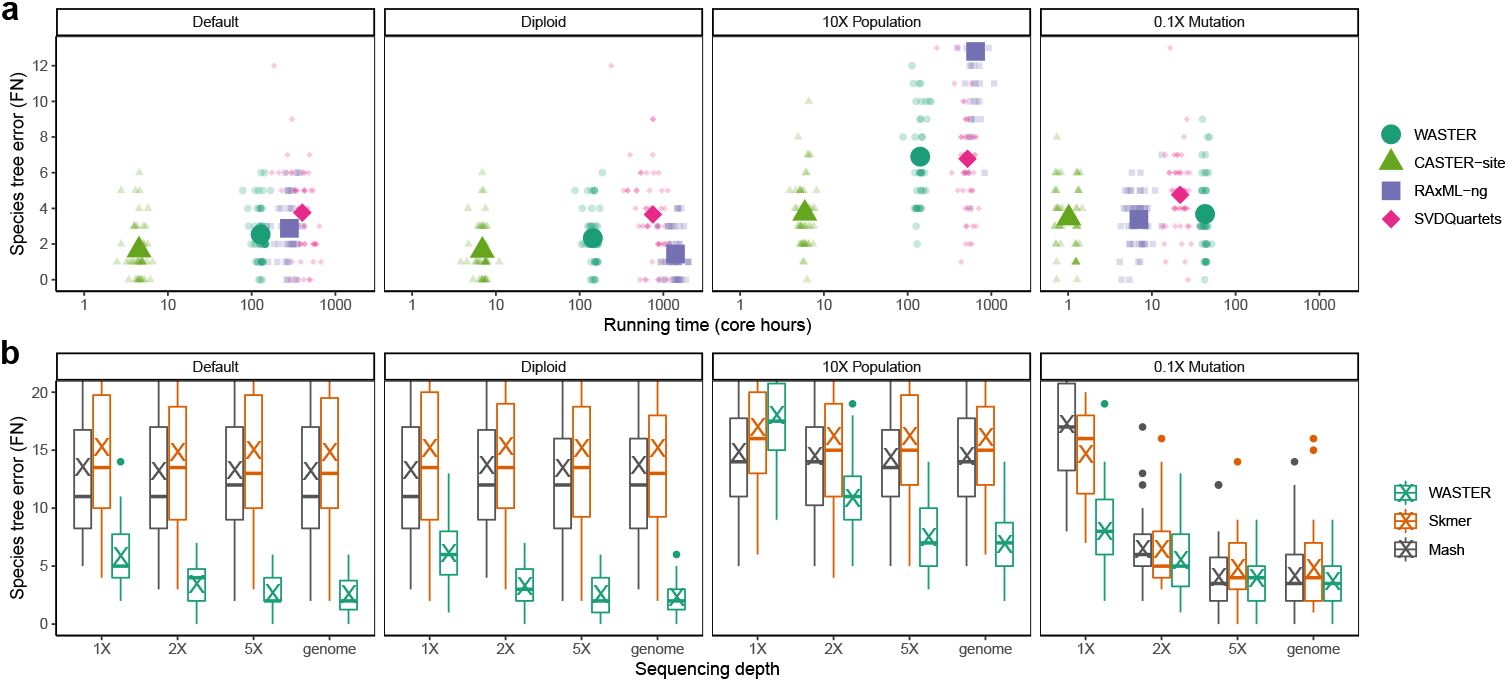
Benchmarking WASTER via R201 simulations. All simulations are performed on the 201-species phylogeny described in [13]. **a**, The number of branches differing between the inferred and true species trees (bipartition false negatives) versus running time (log scale) for WASTER and three alignment-based species tree inference methods using aligned genomes under various settings. Small symbols represent individual replicates, while large, opaque symbols denote mean values across 50 replicates. **b**, The number of branches differing between inferred and true species trees (FN) versus sequencing depth for WASTER, Mash, and Skmer on simulated short reads under different settings. Inferred species trees using true genomes are included as control groups. Under 0.1X mutation rate setting, Skmer fails to output phylogenies on 36/24/25/21 replicates for 1X/2X/5X/genome cases, respectively.

We also explored WASTER’s performance under various settings, contrasting it with alignment-based methods under different ploidies, ILS levels, and mutation rates. Replacing haploid genomes with diploid genomes did not significantly impact (*p ≈* 0.11) WASTER’s accuracy, mirroring results seen with CASTER-site and SVDQuartets. RAxML-ng, employing a diploid model, improved accuracy at the cost of increased runtime. An increase in effective population size by 10X to model elevated ILS levels notably diminished the accuracy of all methods, including WASTER (*p <* 10^*−*15^). Here, RAxML-ng’s performance was notably poorer than WASTER and the other methods. To simulate less genetic divergence among species, we reduced the mutation rate to 10%, resulting in a statistically significant decrease in accuracy for all methods (*p <* 0.05), attributed to a reduction in informative sites. Under this setting, the runtime of alignment-based methods improved substantially compared to the default setting. However, due to bottlenecks in the variable site identification step, WASTER’s runtime improvement was less pronounced (Fig. 2a).

#### 2.2.2 SR201 simulated short reads

We extended our analysis to assess WASTER’s performance utilizing Illumina-like short reads, generated from the SR201 dataset alignments using the ART simulator [29]. This evaluation focused on the impact of varying coverage levels on WASTER’s accuracy. Our findings indicated a direct correlation between increased coverage and enhanced accuracy of WASTER across all settings. A notable observation was that at 5X coverage, the species tree error for WASTER, based on short reads, averaged 4.2 branches, deteriorating by less than 10% compared to the error obtained using true simulated genomes.

Furthermore, we contrasted WASTER’s accuracy with that of Mash [17] and Skmer [19], two frequently used k-mer-based assembly-free and reference-free pairwise evolutionary distance estimators. The k-mer-based distance methods, commonly employed for phylogenetic reconstruction by providing distance matrices to FastME [30], demonstrated consistent species tree errors of approximately 15 branches across all but 0.1 mutation rate settings, irrespective of coverage level. This is likely attributable to their lack of robustness to high evolutionary distances, where fewer k-mers are shared across species, underscoring the superiority of WASTER.

We also evaluated local bootstrap support on WASTER inferred trees in the same simulated dataset. Aggregating across all experimental conditions, replicates, and species tree branches, 94.4% of correctly inferred branches exhibited bootstrap support of at least 95%, while 94.6% of incorrectly inferred branches fell below this threshold, indicating that a 95% cutoff provides well-calibrated measures of statistical uncertainty. At a more stringent cutoff of 99%, 92.2% of correctly inferred branches achieved support of at least 99%, whereas 98.6% of incorrect branches remained below 99% (Fig. A2).

#### 2.2.3 Quartet simulation

We next reused the simulation procedure described in [13] to generate quartet trees for benchmarking WASTER under challenging conditions involving both high levels of incomplete lineage sorting (ILS) and mutation-rate heterogeneity, which together can induce both “long-branch attraction” and “long-branch repulsion.” We simulated 100 replicate alignments, each comprising 20 million base pairs (Mbp), for each of three possible species-tree topologies. Simulations were conducted under the Hudson coalescent model [24] using `msprime` [31], with varying tree height (*λ* = *{*1, 0.5, 0.2*}*) and internal branch length (*x* = *{*0.001, 0.002, 0.005 *}*). The branch lengths for the baseline condition (*λ* = 1, *x* = 0.001) were chosen to reflect those observed in a ratite dataset [32] (Fig. A3a).

Benchmarking results on simulated alignments show that CASTER-site consistently outperforms all other methods across all tested conditions (Fig. A4a). RAxML-ng exhibits signs of long-branch repulsion in the Farris-zone topology (*BC* | *AD*), particularly when the internal branch is short (Fig. A4b). Both alignment-free methods display a substantial increase in species tree estimation error under deep phylogenies (*λ* = 1). In contrast, under shallow phylogenies (*λ≤* 0.5), WASTER outperforms Mash when internal branches are short (*x ≤* 0.002), although WASTER is more sensitive to shorter alignments (*≤* 2 Mbp). In addition to alignments, we also simulated short-read datasets at varying coverage levels (*{*1, 2, 5*}×*) using ART [29] under shallow phylogenies (*λ ≤* 0.5). Overall, WASTER achieves higher accuracy with increasing coverage, while Mash performs better at very low coverage (Fig. A4c). Notably, Mash suffers from long-branch attraction in deeper phylogenies when the internal branch is short (Fig. A4d).

#### 2.2.4 GDL simulation

To evaluate the performance of WASTER and Mash on multi-copy gene families, we extended the quartet simulations to include gene duplication and loss (GDL). Simulations were conducted under varying GDL rates, with tree height and internal branch length fixed at *λ* = 0.2 and *x* = 0.002, respectively. To enable the emergence of gene superfamilies, duplications and losses were simulated beginning 0.07 substitution units above the root (Fig. A3b). We varied the gene duplication rate across 0, *{*1, 2, 4, 8 *}×*, maintaining a fixed 1:1 ratio between gene duplication and loss rates. For each replicate, we simulated 10,000 gene family trees using SimPhy [33], followed by recombination-free alignments of 100 to 10,000 base pairs (bp) generated with INDELible [34]. Additionally, we simulated short-read sequencing data at varying coverage levels (*{*1, 1.5, 2, 5 *}×*) using ART [29].

Simulation results show that Mash performs better when genomes are short and sequencing coverage is low. In contrast, WASTER outperforms Mash for longer genomes and higher coverage levels. Notably, as genome length increases, the species tree error rate of WASTER approaches 0% regardless of the GDL rate or sequence depth, demonstrating statistical consistency under GDL. In comparison, Mash exhibits a marked increase in error rate as the GDL rate increases. Furthermore, at high GDL levels (8 *×*), Mash’s error rate increases with sequencing depth, indicating a lack of statistical consistency under GDL (Fig. A5).

### 2.3 Applications in real data

#### 2.3.1 Applications to raw reads

We also compared phylogenies estimated using WASTER, Mash, and Skmer from low-coverage short reads, to phylogenies constructed using traditional methods employing high-coverage reads. Our analyses span multiple taxonomic groups:

1. **Rats:** We evaluated 18 rat species, particularly focusing on the *Niviventer* genus [35]. The established phylogeny, derived from 14X coverage reads (approximately 39 Gbps), was obtained using MrBayes [36] (concatenation) and ASTRAL [37], both yielding identical results. We downsampled the reads to 1.5X coverage (3.75 Gbps) for reconstructing phylogenies with WASTER, Mash, and Skmer. The phylogeny reconstructed by WASTER matched the established one, while the phylogenies by Mash and Skmer differed from the established phylogeny by two and five branches, respectively (Fig. 3a and A6a).
2. **Butterflies:** This study involved 35 butterfly species from the Nymphalidae family [38], with a published phylogeny derived from RAxML on genome-wide variable sites. We standardized the downsampled reads to 2 Gbps across species. WASTER’s reconstruction deviated from the published phylogeny by three branches. Notably, the phylogeny observed within the *Kallima* genus aligns with the phylogeny inferred by CASTER-site, suggesting potential introgression [13]. The discrepancies in the deeper splits could potentially be attributed to rapid radiation causing short internal branch lengths associated with high statistical uncertainty [38]. Mash and Skmer, however, differed by 8 and 9 branches, respectively (Fig. 3b and A6b).
3. **Birches:** We analyzed high-coverage RAD sequences (3 Gbps on average) for *Betula utilis* FC and *Betula buggsii* [39] and low-coverage RAD sequences (0.3 Gbps on average) for 20 other birch species [40]. The established phylogeny, inferred using RAxML on aligned orthologous internal transcribed spacer (ITS) sequences, was compared with our reconstructions. WASTER, Mash, and Skmer all placed *B. lenta* as the sister of *B. chichibuensis*, rather than the sister of *B. michauxii*, likely due to introgression (Fig. A7ab), which is confirmed using Patterson’s D-statistic [41] (Fig. A7c). Beside *B. lenta*, the WASTER inferred phylogeny differed from the published phylogeny by one branch, while Mash and Skmer differed by five and seven branches, respectively (Fig. 3c and A6c). Noticeably, Mash misgrouped the two species with high coverage as sister taxa, suggesting a potential bias in the method under conditions of non-uniform coverage.
4. **Lizards:** We conducted a similar analysis on 11 Anolis lizard species [42]. The original phylogeny was based on 9X coverage reads. ExaML [43] (concatenation) and ASTRAL generated consistent phylogenies. Upon downsampling to 1.5X coverage (2.25 Gbps), the reconstructed phylogenies by WASTER and Mash matched the published one, while Skmer’s reconstruction deviated by one branch (Fig. A6d).
5. **Mandarin fishes:** Our analysis included three mandarin fish species and one out-group [44], with *Siniperca chuatsi* sequenced using high-coverage long reads and the others with approximately 40X coverage short reads. The published phylogeny was deduced using RAxML [45]. Using downsampled short reads with 2X coverage (1.5 Gbps), as well as the assembled *Siniperca chuatsi* genome, all three methods successfully reconstructed the published phylogeny (Fig. A6e).
6. **Mushrooms:** We examined short reads from four species in the *Tricholomopsis* genus and one outgroup [46]. The established phylogeny, inferred using RAxML on 450 single-copy orthologous genes, was compared with our phylogenies using downsampled reads (1 Gbps). WASTER and Mash successfully reconstructed the published phylogeny, but Skmer did not (Fig. A6f).

**Fig. 3.**
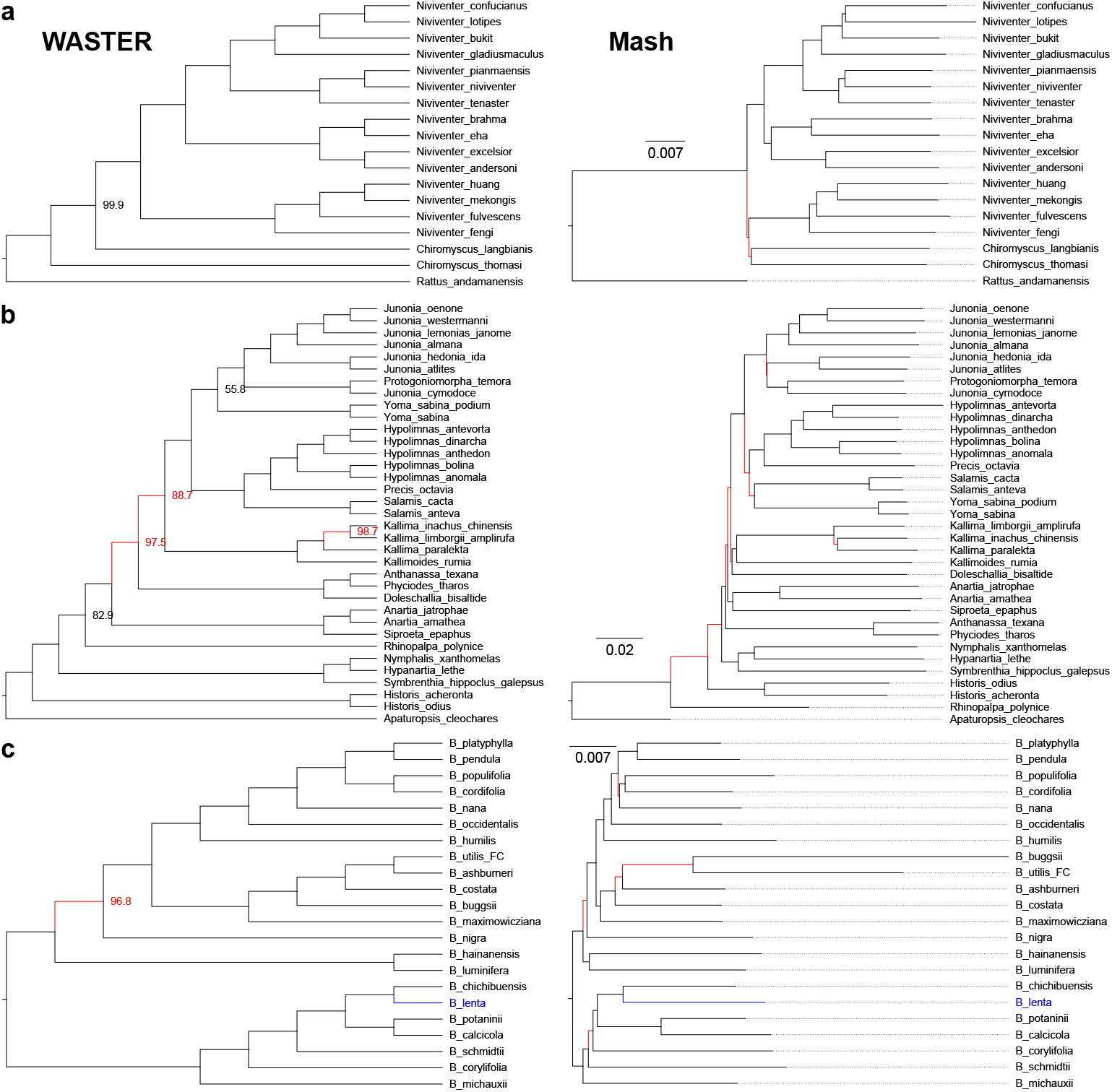
WASTER and Mash on low-coverage short reads. Phylogenies reconstructed using **(a)** 1.5X WGS reads of rat species, (**b**) 2 Gbp butterfly WGS reads, and (**c**) 0.3–3 Gbp birch RAD-seq reads. WASTER branch supports (%) are shown next to the branches, and 100% supports are omitted. Branches that are incongruent with published phylogenies based on high-coverage reads using traditional pipelines are colored red. B. lenta is placed differently from the published phylogeny due to introgression. See also Fig. A6.

These results demonstrate that phylogenies reconstructed using WASTER on low-coverage reads closely align with those derived from traditional pipelines from high-coverage reads, significantly outperforming Mash and, even more so, Skmer in accuracy.

#### 2.3.2 Applications to assembled genomes

In addition to inferring phylogenies of closely related species, WASTER can also generate guide trees for more distantly related species with sufficient accuracy, particularly beneficial when using Progressive Cactus. We demonstrate this with guide trees created by WASTER, Mash, and Skmer for assembled genomes of species in the Primates and Passeriformes orders:

1. **Primates:** Using a dataset of 28 assembled primate genomes [47], WASTER accurately recovered the published phylogeny reconstructed by ExaML on aligned genomes. However, the reconstructions by Mash and Skmer deviated by one and two branches, respectively (Fig. 4a and A8a).
2. **Perching birds:** The dataset of 173 perching bird genomes [10], well-known for their challenging phylogeny, was analyzed. The published phylogeny, reconstructed by ASTRAL+RAxML on aligned genomes, was compared with our reconstructions. WASTER and Mash successfully recovered major clades of Passeriformes (Passerida, Muscicapida, Sylviida, Corvides, and Tyranni), differing from the published phylogeny by 17 and 24 branches, respectively (Fig. 4b). Skmer, however, failed to recover 64 out of 170 branches in the published phylogeny, including misplacing the cinnamon ibon (Hypocryptadius cinnamomeus) among Sylviida, instead of Passerida (Fig. A8b).

**Fig. 4.**
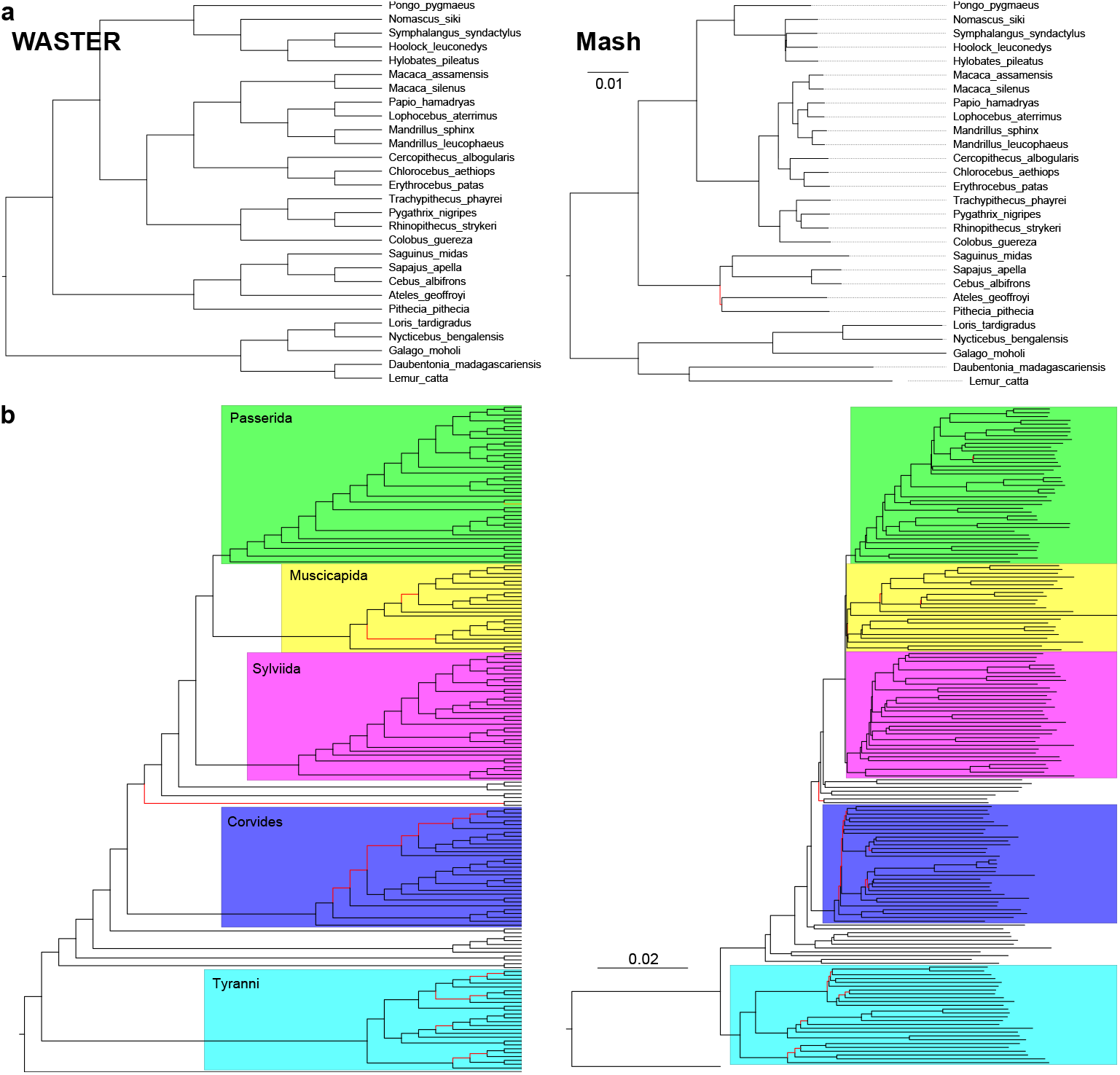
WASTER and Mash on assembled genomes. **a**, Phylogenies reconstructed using 28 assembled primate genomes. WASTER provides 100% supports for all branches. **b**, Phylogenies reconstructed from assembled genomes of 173 perching birds, with species names and support values omitted. Branches incongruent with the published phylogenies are highlighted red. See also Fig. A8.

## 3 Discussion

The increasing availability of whole-genome sequencing data has not resulted in a corresponding increase in phylogenetic inference using aligned whole genomes. A major impediment has been the extensive computational resources and significant human effort required, rendering such approaches impractical for many researchers. WASTER addresses this challenge by offering a computationally efficient and automated solution, facilitating broader adoption of whole-genome phylogenetic analyses. Additionally, WASTER’s output can reduce the cost of many projects by providing high-quality phylogenies using assembly-free low-coverage sequencing, thereby greatly reducing the cost of phylogenomics. This type of application will enable whole genome approaches to be adopted by a much larger share of the taxonomic community that does not have access to the type of resources necessary to produce high-quality assemblies for all species of interest. Even when assembly is possible, WASTER can play a crucial role in verifying the accuracy of the phylogenetic inferences that rely on many steps of assembly and alignment subject to potential human errors. Furthermore, WASTER can be used to produce guide trees for progressive alignment.

WASTER also presents several avenues for future development and refinement. For example, there exists a promising potential for adapting WASTER to identify possible gene-flow events, including ancient ones, using phylogenetic asymmetries and for inferring phylogenetic networks. Another key area for advancement is to extend WASTER to estimate branch lengths in presence of ILS and, potentially, gene-flow. Finally, integrating WASTER with other variable-site-based tools would create a more comprehensive and user-friendly platform, broadening the scope of assembly-free phylogenomic and evolutionary studies.

Despite its broad applicability, WASTER has several limitations:

1. As a k-mer-based method, WASTER is not well suited for reconstructing deep phylogenies, such as intraordinal, family-level insect trees. In such cases, it may produce incorrect or even misleading topologies.

2. Being assembly-free, WASTER assumes genomes lack strand bias and exhibit uniform GC content. In reality, GC content often varies across regions, and strand biases can occur, leading to potential inference errors.

3. Unlike distance-based approaches, WASTER does not provide branch lengths by default. Estimating branch lengths under ILS is already challenging, and ascertainment bias toward conserved regions further complicates this task.

Future work could address these issues by integrating a robust branch length estimation algorithm into WASTER and enhancing its performance on deep phylogenies.

## 4 Methods

### 4.1 WASTER

WASTER initially identifies aligned variable sites using Algorithm 1, described below, before leveraging CASTER-site for species tree inference. Acknowledging its limitation in distinguishing between forward and reverse DNA strands, WASTER operates under the assumption of no strand-bias, equating the equilibrium frequencies of A to T and C to G. Consequently, differently from the conventional CASTER-site that assumes the F84 model, WASTER operates under the T92 model. This model can be easily reduced from F84 model by assuming equal equilibrium frequencies of complementary nucleotides (Fig. 5). WASTER computes local bootstrap support for each branch using 1,000 site-based bootstrap replicates. The local support for a branch is defined as the percentage of replicates in which the species tree topology is favored over both of its nearest-neighbor interchange (NNI) alternatives around that branch.

**Fig. 5.**
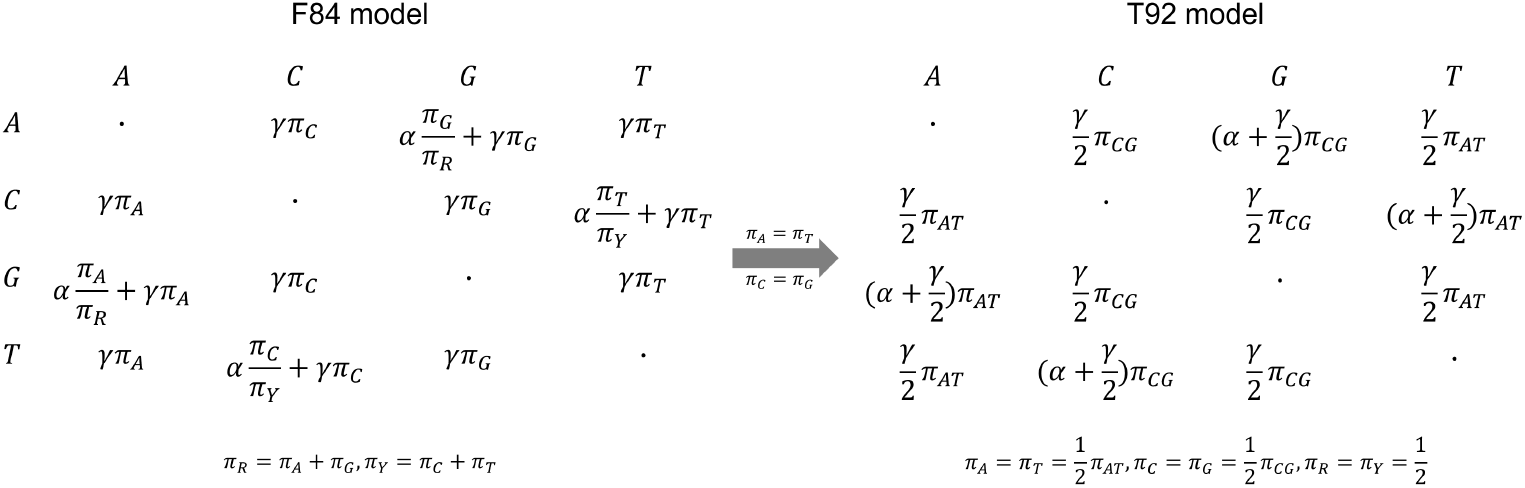
No-strand bias model. By assuming equal equilibrium frequencies of complementary nucleotides, the F84 model reduces to the T92 model.

### 4.2 Simulating low-coverage reads

Reads with 1X, 2X, and 5X coverage in the simulation studies were simulated using ART Illumina v2.5.8 with the following command:

#### Algorithm 1

WASTER’s algorithm to indentify a set of aligned variable sites *S* using sets of k-mers *K*_1_, …, *K*_*n*_, where each set represents all k-mers in the input reads/genomes of a species.

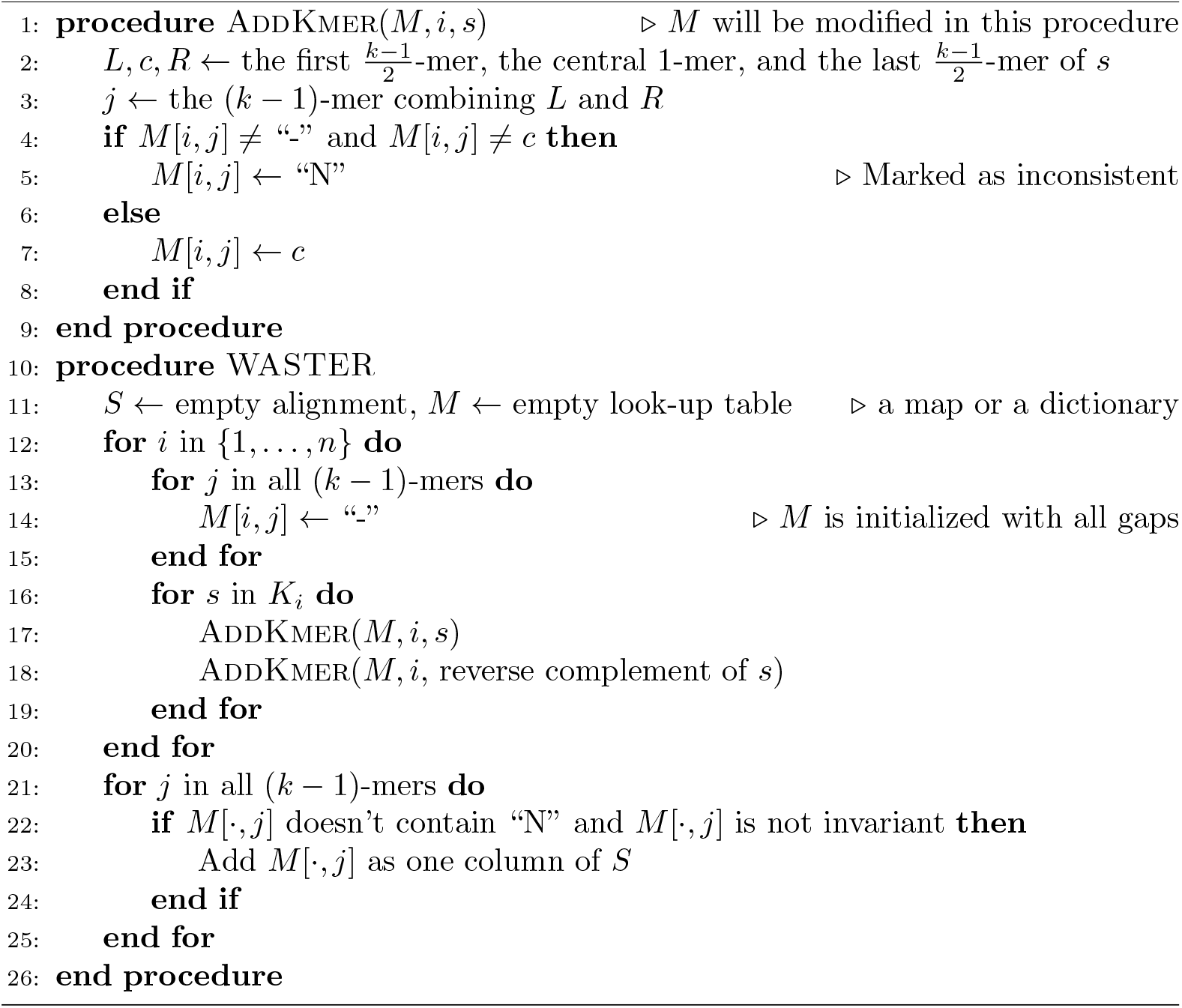

for i in 1 2 5 ; do

art_illumina –ss HS25 –sam -i $GENOME –p –l 150 –f $ i \

–m 800 –s 20 –o ${i}X_READS

cat ${i}X_READS{1,2}. fq > ${i}X_READS_COMBINED. fq

done

Each input file contains one FASTA sequence representing a haploid genome or two FASTA sequences representing a diploid genome. Each output file contains reads from both ends.

### 4.3 Species tree inference

We inferred species tree from simulated and real data using WASTER, Mash, and Skmer. Aside from those alignment free methods, we also reuse inferred species trees by CASTER-site, RAxML-ng, and SVDQuartets on simulated true alignments from a previous study [13] for comparison (Fig. 2a).

#### 4.3.1 WASTER

The following command was used to infer species trees:

waster −C −t 16 −i $INPUT −o $OUTPUT

The input file here is a list of species names followed by the corresponding input files:

SpeciesName1 Species 1.fq

SpeciesName2 Species 2.fq

…

#### 4.3.2 Mash

The following script was used to infer species trees:

mash triangle −s 1000000 $INPUT *>* mash . tsv

python3 mashFormatFastme.py mash.tsv mash.phy

fastme −i mash.phy −o $OUTPUT

The python script “mashFormatFastme.py” was used to format the output distance matrix by Mash as the input format of FastME:

**import** sys

with **open** (sys.argv[2], “w”) as f :

lines = []

**for** line **in open** (sys . argv [1]) :

lines . append (line.split ())

n = **int** (lines [0] [0])

f.write (lines [0] [0] + “ *\*n”)

**for** i **in range** (1, n+1):

f.write (“ *\* t “ .join ([lines [i] [0] . split (“ . “)[0]]

+ [lines[i] [j] **for** j **in range** (1,i)] + [“0 “]

+ [lines[j] [i] **for** j **in range** (i +1, n +1)]) + “*\*n”)

#### 4.3.3 Skmer

The following script was used to infer species trees:

skmer reference $INPUT −p 16 −t #generating ref −dist−mat.txt bash skmerFormatFastme.bash ref −dist −mat.txt skmer.phy

fastme −i skmer.phy −o $OUTPUT

the python script “skmerFormatFastme.bash” was used to format the output distance matrix by Skmer as the input format of FastME:

tail −n +2 $1 |wc −l *>* $2

tail −n +2 $1 *>>* $2

**sed** −i “ s /*\* t / - - / g” $2

## Acknowledgments

This work was supported by a research grant (40582) from VILLUM FONDEN and NIH grant R35 GM153400-01. We thank Yulong Xie from Zhejiang university for advise on avian phylogenetics.

## Appendix A Supplementary figures

**Fig. A1.**
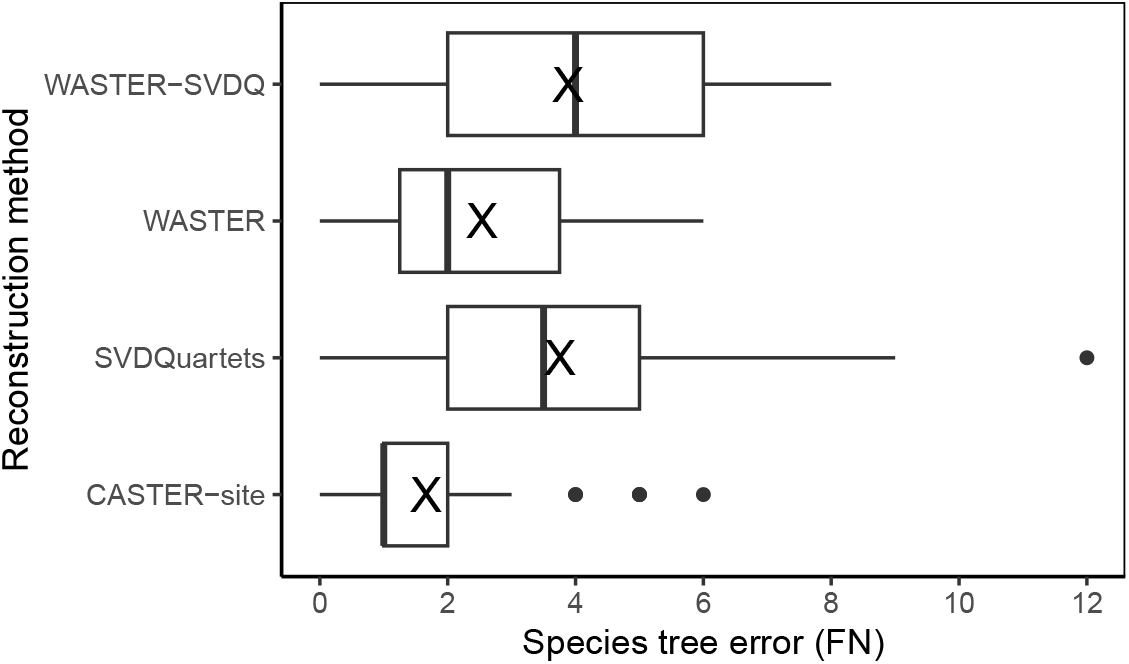
CASTER-site versus SVDQuartets on variable sites extracted by WASTER. Species trees were inferred using SVDQuartets applied to variable sites extracted by WASTER (denoted WASTER-SVDQ) on 25 replicate alignments under the default condition of the SR201 dataset. The resulting species tree errors were compared against those of standard WASTER, standalone SVDQuartets, and CASTER. WASTER outperformed WASTER-SVDQ, indicating that its integrated use of CASTER-site is more effective than coupling with SVDQuartets.

**Fig. A2.**
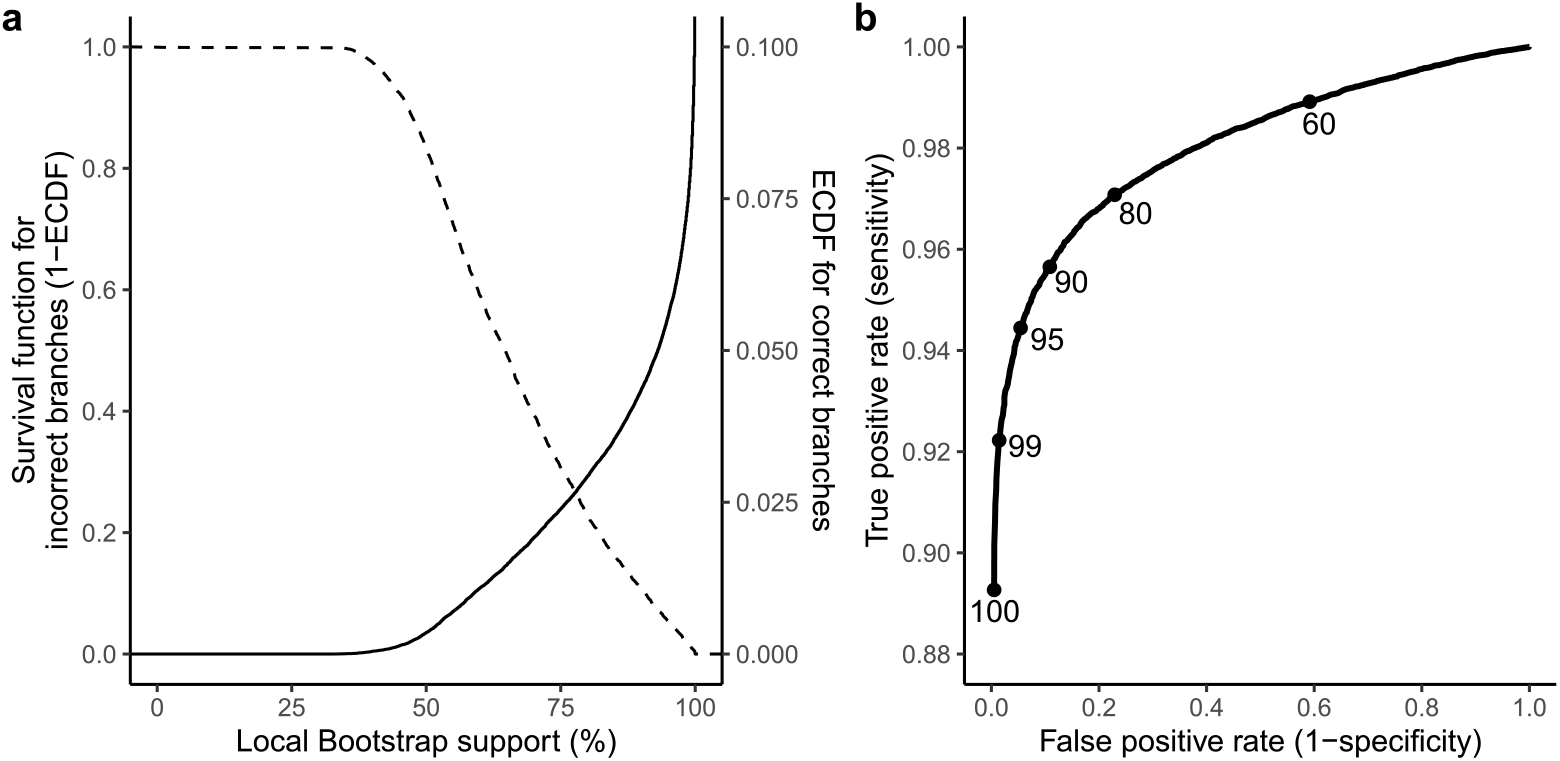
Evaluation of local bootstrap support on SR201 dataset. (**a**) Empirical cumulative distribution function (ECDF) of support values for correctly inferred branches (solid line) and survival function (1-ECDF) for incorrect branches (dashed line). (**b**) Receiver operating characteristic (ROC) curves of support values. True positive rate (sensitivity) and false positive rate (1-specificity) at *{*60,80,90,95,99,100*}*% supports are highlighted as dots.

**Fig. A3.**
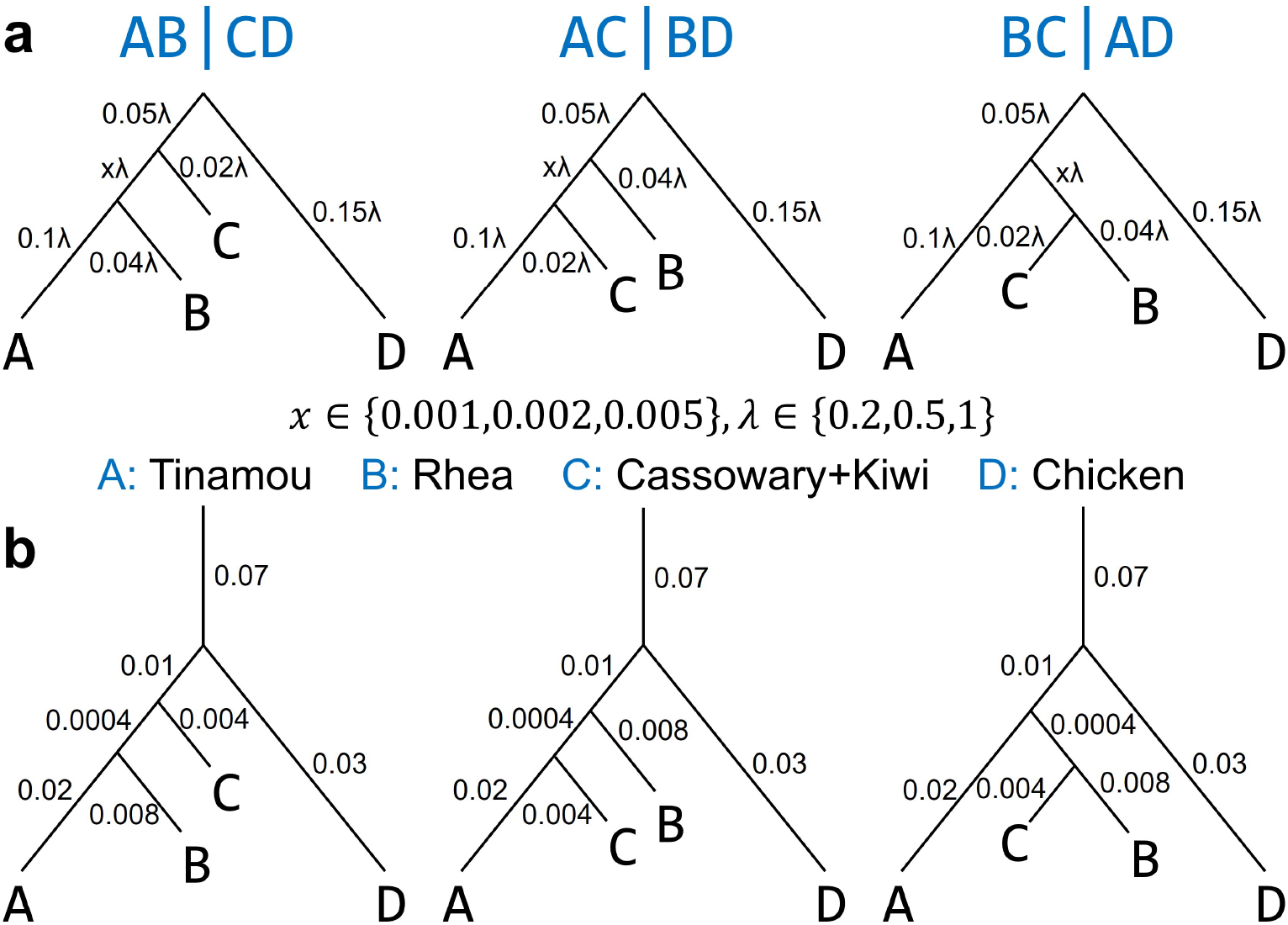
Experimental setup of additional simulation studies. (**a**) Quartet dataset. Aligned genomes are simulated from three topologies corresponding to Felsenstein’s (*AB* | *CD* and *AC* | *BD*) and Farris’s zone (*BC* | *AD*) under various tree height (*λ*) and internal branch length (*x*). The branch lengths are labelled in substitution units. In all conditions, a consistent level of ILS is maintained by fixing the discordance between the species tree and the local (gene) trees at 60%. (**b**) GDL dataset. In this dataset, tree height (*λ* = 0.2) and internal branch length (*x* = 0.002) are fixed. Gene duplication and loss simulation starts 0.07 substitution unit above the root.

**Fig. A4.**
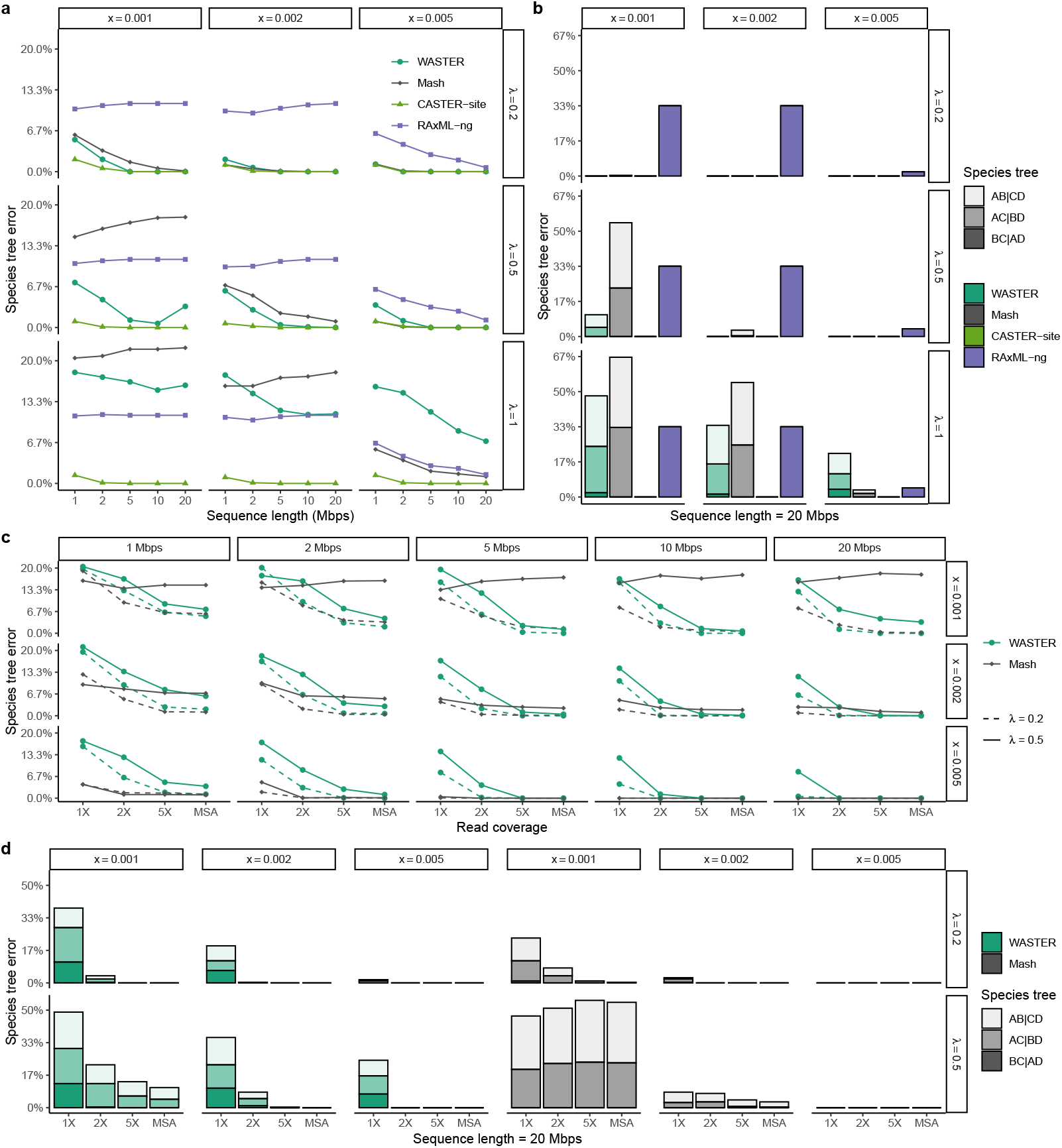
Benchmarking via Quartet dataset. Species tree estimation error of alignment-free and alignment-based methods on true MSAs by (**a**) sequence length and (**b**) true species tree topology. Species tree estimation error of assembly-free methods on simulated short reads and true MSAs by (**c**) sequence length and (**d**) true species tree topology under shallow phylogenies (*λ ≤* 0.5).

**Fig. A5.**
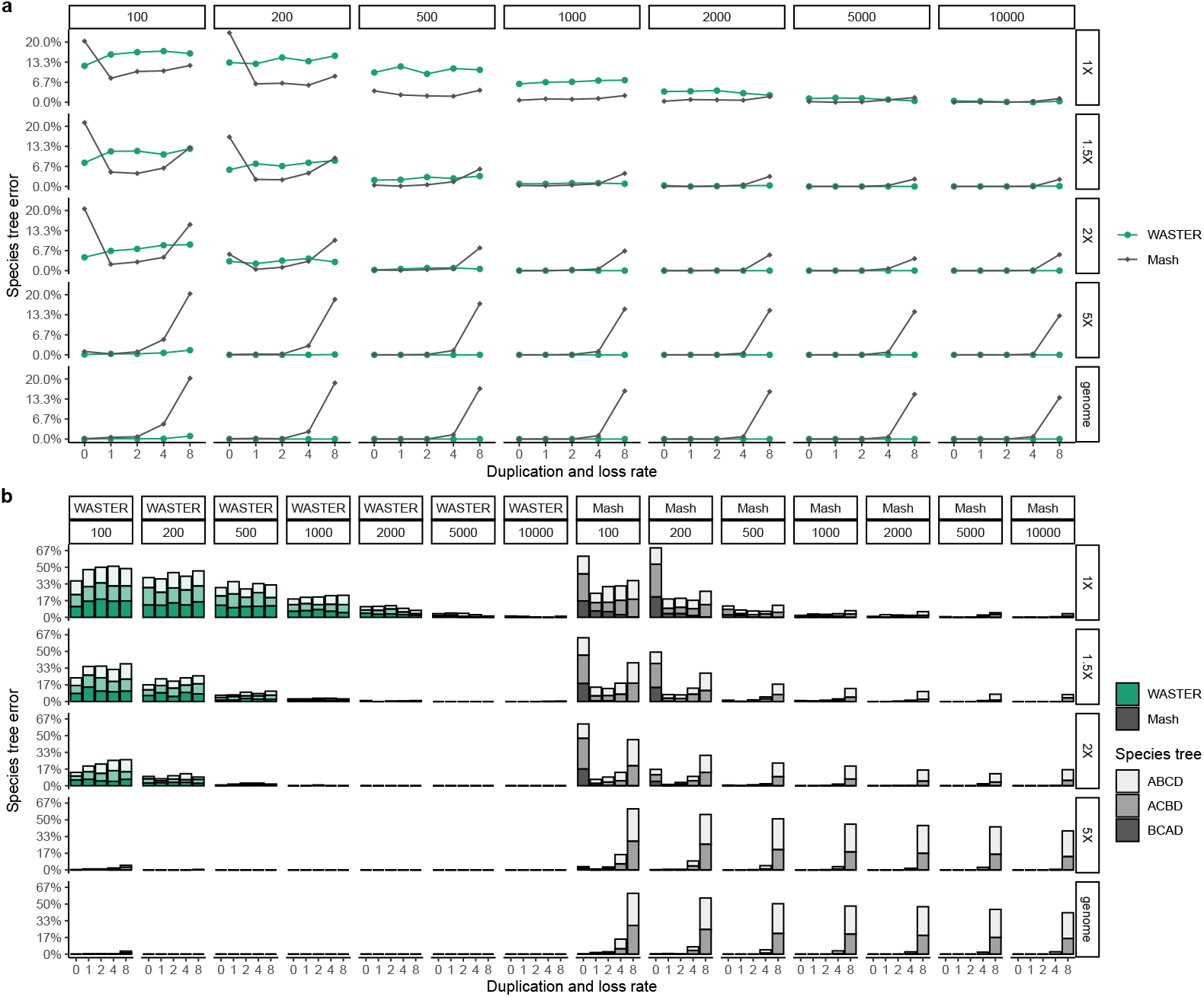
Benchmarking via GDL dataset. Species tree estimation error of WASTER and Mash on simulated short reads and true MSAs across (**a**) gene duplication and loss rates and (**b**) duploss rates and true species tree topology under various sequencing depths (rows) and gene lengths (columns).

**Fig. A6.**
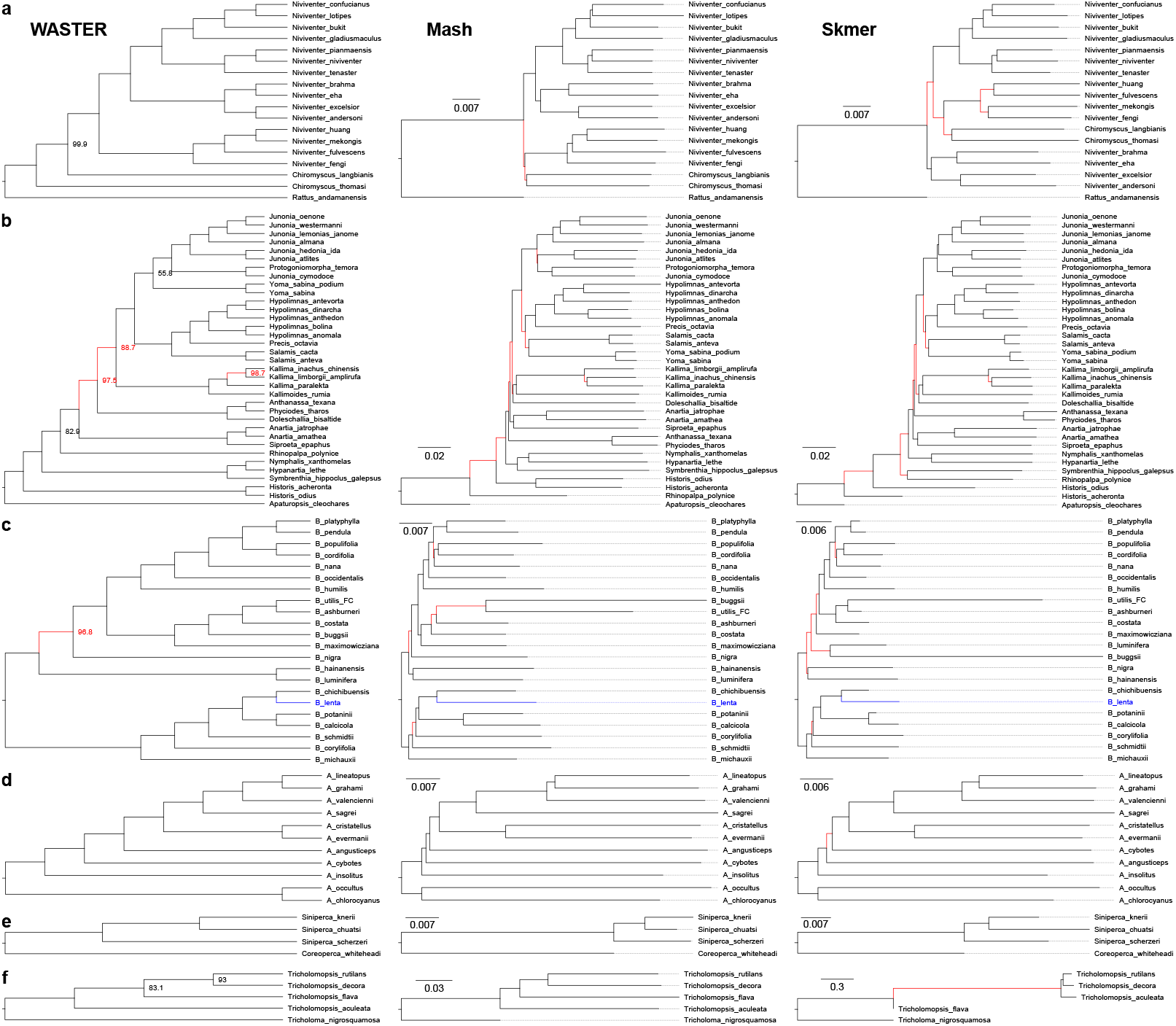
WASTER, Mash, and Skmer on low-coverage short reads. Phylogenies reconstructed using (**a**) 1.5X WGS reads of rat species, (**b**) 2Gbp butterfly WGS reads, (**c**) 0.3-3Gbp birch RAD reads, (**d**) 1.5X lizard WGS reads, and (**e**) 1.5Gbp mandarin fish WGS reads, and (**f**) 1Gbp mushroom WGS reads. WASTER branch supports (%) are shown next to the branches, and 100% supports are omitted. Branches that are incongruent with published phylogenies based on high-coverage reads using traditional pipelines are colored red. B. lenta is placed differently from the published phylogeny due to introgression.

**Fig. A7.**
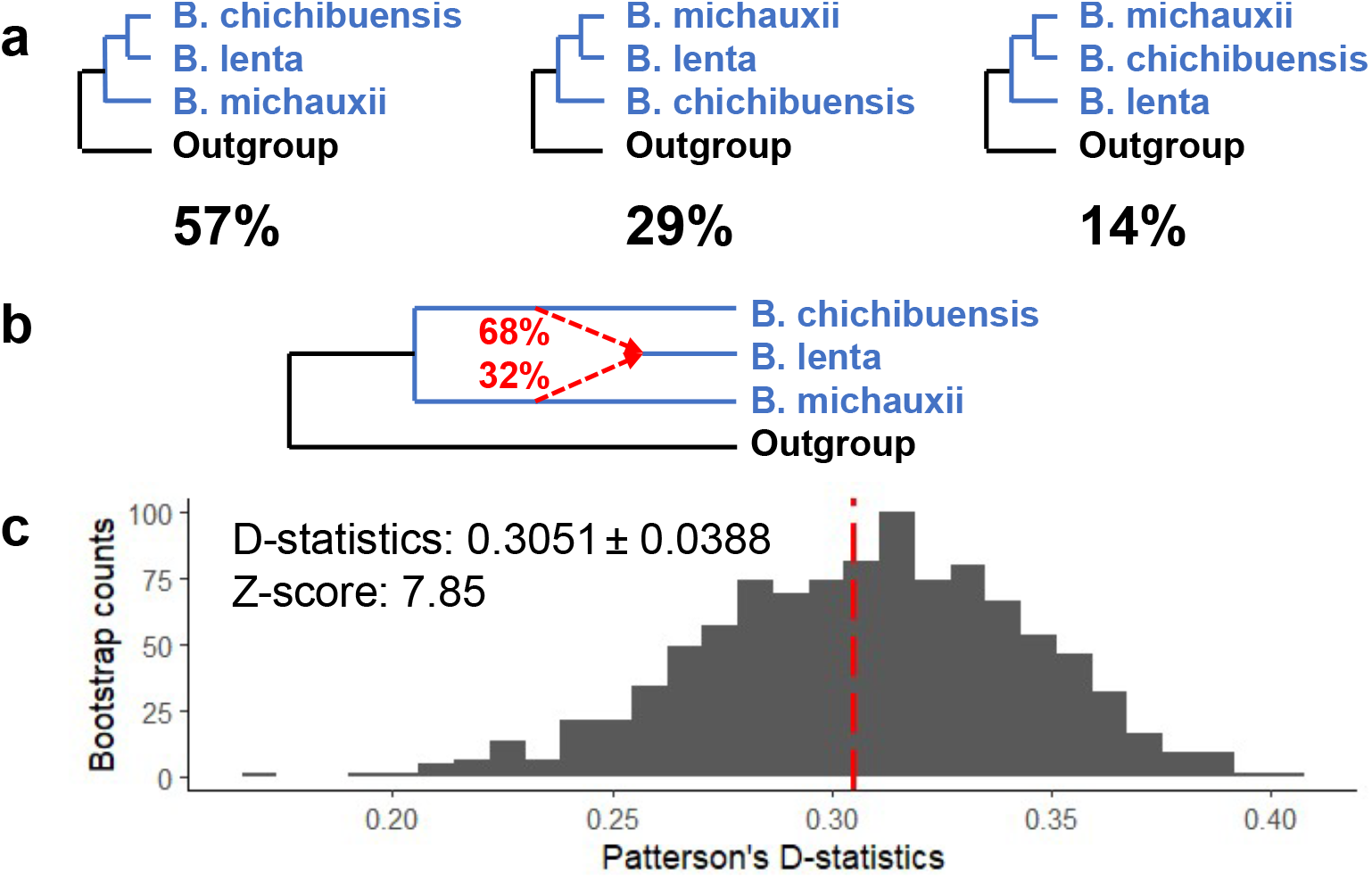
Non-ILS-like signal around B. lenta. **a**, Normalized WASTER scores for all quartet topologies with species B. chichibuensis, B. lenta, B. michauxii, and B. maximowicziana (outgroup). Notably, in the absence of introgression, the second and third topologies would be expected to have similar WASTER scores. **b**, One plausible explanation for the observed disparity in WASTER scores is hybrid speciation. In this scenario, 68% of B. lenta genome is derived from B. chichibuensis and 32% originates from B. michauxii. **c**, A histogram of Patterson’s D-statistic on 1,000 block bootstrap replicates confirms a non-ILS-like signal with 100% bootstrap support. The D-statistic is based on 4,666 alignments of RAD sequences and the species tree topology (((B. chichibuensis, B. lenta), B. michauxii), Outgroup). The dashed line represents the D-statistic of 0.3051 without bootstrapping. The standard deviation based on 1,000 bootstraps is 0.0388, yielding a Z-value of 7.85.

**Fig. A8.**
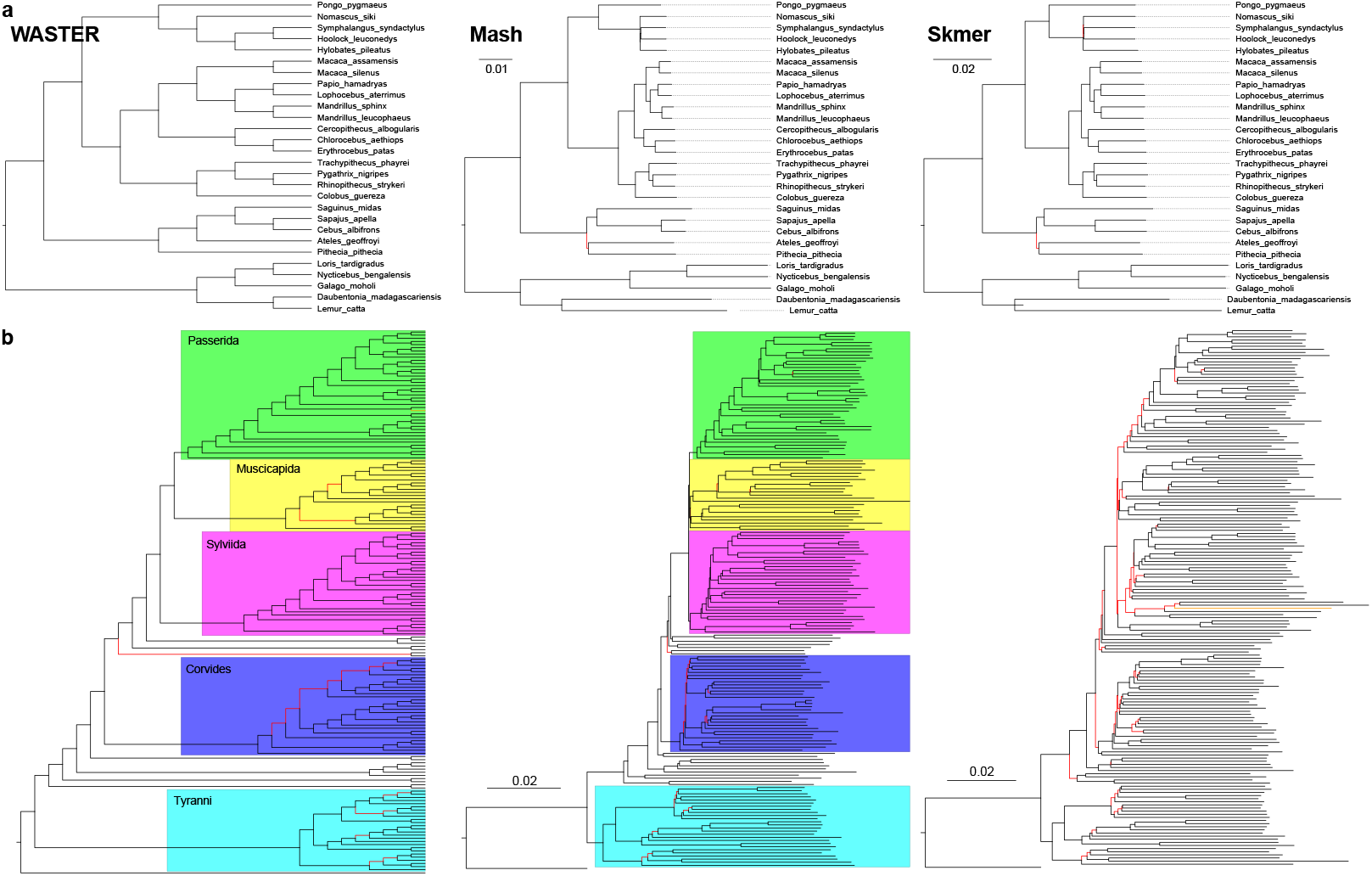
WASTER, Mash, and Skmer on assembled genomes. **a**, Phylogenies reconstructed using 28 assembled primate genomes. **b**, Phylogenies reconstructed using assembled genomes of 173 perching birds, with species names and support values omitted. The terminal branch corresponding to cinnamon ibon (Hypocryptadius cinnamomeus) is colored orange. Branches incongruent with the published phylogenies are highlighted red.

## References

[1] Lewin, H.A., Robinson, G.E., Kress, W.J., Baker, W.J., Coddington, J., Crandall, K.A., Durbin, R., Edwards, S.V., Forest, F., Gilbert, M.T.P., Goldstein, M.M., Grigoriev, I.V., Hackett, K.J., Haussler, D., Jarvis, E.D., Johnson, W.E., Patrinos, A., Richards, S., Castilla-Rubio, J.C., Van Sluys, M.A., Soltis, P.S., Xu, X., Yang, H., Zhang, G.: Earth BioGenome Project: Sequencing life for the future of life. Proceedings of the National Academy of Sciences of the United States of America 115(17), 4325–4333 (2018). 10.1073/PNAS.1720115115/SUPPL{}FILE/PNAS.1720115115.SAPP.

[2] Haussler, D., O’Brien, S.J., Ryder, O.A., Keith Barker, F., Clamp, M., Crawford, A.J., Hanner, R., Hanotte, O., Johnson, W.E., McGuire, J.A., Miller, W., Murphy, R.W., Murphy, W.J., Sheldon, F.H., Sinervo, B., Venkatesh, B., Wiley, E.O., Allendorf, F.W., Amato, G., Scott Baker, C., Bauer, A., Beja-Pereira, A., Bermingham, E., Bernardi, G., Bonvicino, C.R., Brenner, S., Burke, T., Cracraft, J., Diekhans, M., Edwards, S., Ericson, P.G.P., Estes, J., Fjelsda, J., Flesness, N., Gamble, T., Gaubert, P., Graphodatsky, A., Marshall Graves, J.A., Green, E.D., Green, R.E., Hackett, S., Hebert, P., Helgen, K.M., Joseph, L., Kessing, B., Kingsley, D.M., Lewin, H.A., Luikart, G., Martelli, P., Moreira, M.A.M., Nguyen, N., Ortí, G., Pike, B.L., Rawson, D.M., Schuster, S.C., Seuánez, H.N., Bradley Shaffer, H., Springer, M.S., Stuart, J.M., Sumner, J., Teeling, E., Vrijenhoek, R.C., Ward, R.D., Warren, W.C., Wayne, R., Williams, T.M., Wolfe, N.D., Zhang, Y.P., Graves, J., Springer, M., Williams, T., Wolfe, N., Edwards, S.V., Orti, G., Rawson, D.M., Felsenfeld, A., Seuanez, H.N., Stuart, J.M., Turner, S.: Genome 10K: A Proposal to Obtain Whole-Genome Sequence for 10000 Vertebrate Species. Journal of Heredity 100(6), 659–674 (2009). 10.1093/JHERED/ESP086

[3] Zhang, G.: Bird sequencing project takes off. Nature 2015 522:7554 522(7554), 34–34 (2015). 10.1038/522034d

[4] Fan, G., Song, Y., Yang, L., Huang, X., Zhang, S., Zhang, M., Yang, X., Chang, Y., Zhang, H., Li, Y., Liu, S., Yu, L., Chu, J., Seim, I., Feng, C., Near, T.J., Wing, R.A., Wang, W., Kun Wan, K., Wang, J., Xu, X., Yang, H., Liu, X., Chen, N., He, S.: Initial data release and announcement of the 10,000 Fish Genomes Project (Fish10K). GigaScience 9(8) (2020). 10.1093/GIGASCIENCEGIAA080

[5] Evans, J.D., Brown, S.J., Hackett, K.J.J., Robinson, G., Richards, S., Lawson, D., Elsik, C., Coddington, J., Edwards, O., Emrich, S., Gabaldon, T., Goldsmith, M., Hanes, G., Misof, B., Muñoz-Torres, M., Niehuis, O., Papanicolaou, A., Pfrender, M., Poelchau, M., Purcell-Miramontes, M., Robertson, H.M., Ryder, O., Tagu, D., Torres, T., Zdobnov, E., Zhang, G., Zhou, X.: The i5K Initiative: Advancing Arthropod Genomics for Knowledge, Human Health, Agriculture, and the Environment. Journal of Heredity 104(5), 595–600 (2013). 10.1093/JHERED/EST050

[6] GIGA Community of Scientists: The Global Invertebrate Genomics Alliance (GIGA): Developing Community Resources to Study Diverse Invertebrate Genomes. Journal of Heredity 105(1), 1–18 (2014). 10.1093/JHERED/EST084

[7] Cheng, S., Melkonian, M., Smith, S.A., Brockington, S., Archibald, J.M., Delaux, P.M., Li, F.W., Melkonian, B., Mavrodiev, E.V., Sun, W., Fu, Y., Yang, H., Soltis, D.E., Graham, S.W., Soltis, P.S., Liu, X., Xu, X., Wong, G.K.S.: 10KP: A phylodiverse genome sequencing plan. GigaScience 7(3), 1–9 (2018). 10.1093/GIGASCIENCE/GIY013

[8] Grigoriev, I.V., Nikitin, R., Haridas, S., Kuo, A., Ohm, R., Otillar, R., Riley, R., Salamov, A., Zhao, X., Korzeniewski, F., Smirnova, T., Nordberg, H., Dubchak, I., Shabalov, I.: MycoCosm portal: gearing up for 1000 fungal genomes. Nucleic Acids Research 42(D1), 699–704 (2014). 10.1093/NAR/GKT1183

[9] Foley, N.M., Mason, V.C., Harris, A.J., Bredemeyer, K.R., Damas, J., Lewin, H.A., Eizirik, E., Gatesy, J., Karlsson, E.K., Lindblad-Toh, K., Springer, M.S., Murphy, W.J., Andrews, G., Armstrong, J.C., Bianchi, M., Birren, B.W., Bredemeyer, K.R., Breit, A.M., Christmas, M.J., Clawson, H., Damas, J., Di Palma, F., Diekhans, M., Dong, M.X., Eizirik, E., Fan, K., Fanter, C., Foley, N.M., Forsberg-Nilsson, K., Garcia, C.J., Gatesy, J., Gazal, S., Genereux, D.P., Goodman, L., Grimshaw, J., Halsey, M.K., Harris, A.J., Hickey, G., Hiller, M., Hindle, A.G., Hubley, R.M., Hughes, G.M., Johnson, J., Juan, D., Kaplow, I.M., Karlsson, E.K., Keough, K.C., Kirilenko, B., Koepfli, K.-P., Korstian, J.M., Kowalczyk, A., Kozyrev, S.V., Lawler, A.J., Lawless, C., Lehmann, T., Levesque, D.L., Lewin, H.A., Li, X., Lind, A., Lindblad-Toh, K., Mackay-Smith, A., Marinescu, V.D., Marques-Bonet, T., Mason, V.C., Meadows, J.R.S., Meyer, W.K., Moore, J.E., Moreira, L.R., Moreno-Santillan, D.D., Morrill, K.M., Muntané, G., Murphy, W.J., Navarro, A., Nweeia, M., Ortmann, S., Osmanski, A., Paten, B., Paulat, N.S., Pfenning, A.R., Phan, B.N., Pollard, K.S., Pratt, H.E., Ray, D.A., Reilly, S.K., Rosen, J.R., Ruf, I., Ryan, L., Ryder, O.A., Sabeti, P.C., Schäffer, D.E., Serres, A., Shapiro, B., Smit, A.F.A., Springer, M., Srinivasan, C., Steiner, C., Storer, J.M., Sullivan, K.A.M., Sullivan, P.F., Sundström, E., Supple, M.A., Swofford, R., Talbot, J.-E., Teeling, E., Turner-Maier, J., Valenzuela, A., Wagner, F., Wallerman, O., Wang, C., Wang, J., Weng, Z., Wilder, A.P., Wirthlin, M.E., Xue, J.R., Zhang, X.: A genomic timescale for placental mammal evolution. Science 380(6643) (2023). 10.1126/SCIENCE.ABL8189

[10] Stiller, J., Feng, S., Chowdhury, A.-A., Rivas-González, I., Duchêne, D.A., Fang, Q., Deng, Y., Kozlov, A., Stamatakis, A., Claramunt, S., Nguyen, J.M.T., Ho, S.Y.W., Faircloth, B.C., Haag, J., Houde, P., Cracraft, J., Balaban, M., Mai, U., Chen, G., Gao, R., Zhou, C., Xie, Y., Huang, Z., Cao, Z., Yan, Z., Ogilvie, H.A., Nakhleh, L., Lindow, B., Morel, B., Fjeldså, J., Hosner, P.A., da Fonseca, R.R., Petersen, B., Tobias, J.A., Székely, T., Kennedy, J.D., Reeve, A.H., Liker, A., Stervander, M., Antunes, A., Tietze, D.T., Bertelsen, M.F., Lei, F., Rahbek, C., Graves, G.R., Schierup, M.H., Warnow, T., Braun, E.L., Gilbert, M.T.P., Jarvis, E.D., Mirarab, S., Zhang, G.: Complexity of avian evolution revealed by family-level genomes. Nature 2024 629:8013 629(8013), 851–860 (2024). 10.1038/s41586-024-07323-1

[11] Kapli, P., Yang, Z., Telford, M.J.: Phylogenetic tree building in the genomic age. Nature Reviews Genetics 2020 21:7 21(7), 428–444 (2020). 10.1038/s41576-020-0233-0

[12] Liu, L., Zhang, J., Rheindt, F.E., Lei, F., Qu, Y., Wang, Y., Zhang, Y., Sullivan, C., Nie, W., Wang, J., Yang, F., Chen, J., Edwards, S.V., Meng, J., Wu, S.: Genomic evidence reveals a radiation of placental mammals uninterrupted by the KPg boundary. Proceedings of the National Academy of Sciences of the United States of America 114(35), 7282–7290 (2017). 10.1073/PNAS.1616744114/SUPPL{}FILE/PNAS.1616744114.SD11.

[13] Zhang, C., Nielsen, R., Mirarab, S.: CASTER: Direct species tree inference from whole-genome alignments. Science 387(6737) (2025). 10.1126/SCIENCE.ADK9688

[14] Genereux, D.P., Serres, A., Armstrong, J., Johnson, J., Marinescu, V.D., Murén, E., Juan, D., Bejerano, G., Casewell, N.R., Chemnick, L.G., Damas, J., Di Palma, F., Diekhans, M., Fiddes, I.T., Garber, M., Gladyshev, V.N., Goodman, L., Haerty, W., Houck, M.L., Hubley, R., Kivioja, T., Koepfli, K.P., Kuderna, L.F.K., Lander, E.S., Meadows, J.R.S., Murphy, W.J., Nash, W., Noh, H.J., Nweeia, M., Pfenning, A.R., Pollard, K.S., Ray, D.A., Shapiro, B., Smit, A.F.A., Springer, M.S., Steiner, C.C., Swofford, R., Taipale, J., Teeling, E.C., Turner-Maier, J., Alfoldi, J., Birren, B., Ryder, O.A., Lewin, H.A., Paten, B., Marques-Bonet, T., Lindblad-Toh, K., Karlsson, E.K.: A comparative genomics multitool for scientific discovery and conservation. Nature 2020 587:7833 587(7833), 240–245 (2020). 10.1038/s41586-020-2876-6

[15] Feng, S., Stiller, J., Deng, Y., Armstrong, J., Fang, Q., Reeve, A.H., Xie, D., Chen, G., Guo, C., Faircloth, B.C., Petersen, B., Wang, Z., Zhou, Q., Diekhans, M., Chen, W., Andreu-Sánchez, S., Margaryan, A., Howard, J.T., Parent, C., Pacheco, G., Sinding, M.H.S., Puetz, L., Cavill, E., Ribeiro, A.M., Eckhart, L., Fjeldså, J., Hosner, P.A., Brumfield, R.T., Christidis, L., Bertelsen, M.F., Sicheritz-Ponten, T., Tietze, D.T., Robertson, B.C., Song, G., Borgia, G., Claramunt, S., Lovette, I.J., Cowen, S.J., Njoroge, P., Dumbacher, J.P., Ryder, O.A., Fuchs, J., Bunce, M., Burt, D.W., Cracraft, J., Meng, G., Hackett, S.J., Ryan, P.G., Jønsson, K.A., Jamieson, I.G., da Fonseca, R.R., Braun, E.L., Houde, P., Mirarab, S., Suh, A., Hansson, B., Ponnikas, S., Sigeman, H., Stervander, M., Frandsen, P.B., van der Zwan, H., van der Sluis, R., Visser, C., Balakrishnan, C.N., Clark, A.G., Fitzpatrick, J.W., Bowman, R., Chen, N., Cloutier, A., Sackton, T.B., Edwards, S.V., Foote, D.J., Shakya, S.B., Sheldon, F.H., Vignal, A., Soares, A.E.R., Shapiro, B., González-Solís, J., Ferrer-Obiol, J., Rozas, J., Riutort, M., Tigano, A., Friesen, V., Dalén, L., Urrutia, A.O., Székely, T., Liu, Y., Campana, M.G., Corvelo, A., Fleischer, R.C., Rutherford, K.M., Gemmell, N.J., Dussex, N., Mouritsen, H., Thiele, N., Delmore, K., Liedvogel, M., Franke, A., Hoeppner, M.P., Krone, O., Fudickar, A.M., Milá, B., Ketterson, E.D., Fidler, A.E., Friis, G., Parody-Merino, A.M., Battley, P.F., Cox, M.P., Lima, N.C.B., Prosdocimi, F., Parchman, T.L., Schlinger, B.A., Loiselle, B.A., Blake, J.G., Lim, H.C., Day, L.B., Fuxjager, M.J., Baldwin, M.W., Braun, M.J., Wirthlin, M., Dikow, R.B., Ryder, T.B., Camenisch, G., Keller, L.F., DaCosta, J.M., Hauber, M.E., Louder, M.I.M., Witt, C.C., McGuire, J.A., Mudge, J., Megna, L.C., Carling, M.D., Wang, B., Taylor, S.A., Del-Rio, G., Aleixo, A., Vasconcelos, A.T.R., Mello, C.V., Weir, J.T., Haussler, D., Li, Q., Yang, H., Wang, J., Lei, F., Rahbek, C., Gilbert, M.T.P., Graves, G.R., Jarvis, E.D., Paten, B., Zhang, G.: Dense sampling of bird diversity increases power of comparative genomics. Nature 2020 587:7833 587(7833), 252–257 (2020). 10.1038/s41586-020-2873-9

[16] Armstrong, J., Hickey, G., Diekhans, M., Fiddes, I.T., Novak, A.M., Deran, A., Fang, Q., Xie, D., Feng, S., Stiller, J., Genereux, D., Johnson, J., Marinescu, V.D., Alföldi, J., Harris, R.S., Lindblad-Toh, K., Haussler, D., Karlsson, E., Jarvis, E.D., Zhang, G., Paten, B.: Progressive Cactus is a multiple-genome aligner for the thousand-genome era. Nature 2020 587:7833 587(7833), 246–251 (2020). 10.1038/s41586-020-2871-y

[17] Ondov, B.D., Treangen, T.J., Melsted, P., Mallonee, A.B., Bergman, N.H., Koren, S., Phillippy, A.M.: Mash: Fast genome and metagenome distance estimation using MinHash. Genome Biology 17(1), 1–14 (2016). 10.1186/S13059-016-0997-X/FIGURES/5

[18] Jain, C., Rodriguez-R, L.M., Phillippy, A.M., Konstantinidis, K.T., Aluru, S.: High throughput ANI analysis of 90K prokaryotic genomes reveals clear species boundaries. Nature Communications 2018 9:1 9(1), 1–8 (2018). 10.1038/s41467-018-07641-9

[19] Sarmashghi, S., Bohmann, K., Gilbert, P.M.T., Bafna, V., Mirarab, S.: Skmer: Assembly-free and alignment-free sample identification using genome skims. Genome Biology 20(1), 34 (2019). 10.1186/s13059-019-1632-4

[20] Balaban, M., Sarmashghi, S., Mirarab, S.: APPLES: Scalable Distance-Based Phylogenetic Placement with or without Alignments. Systematic Biology 69(3), 566–578 (2020). 10.1093/SYSBIO/SYZ063

[21] Dylus, D., Altenhoff, A., Majidian, S., Sedlazeck, F.J., Dessimoz, C.: Inference of phylogenetic trees directly from raw sequencing reads using Read2Tree. Nature Biotechnology 2023, 1–9 (2023). 10.1038/s41587-023-01753-4

[22] Schwartz, R.S., Harkins, K.M., Stone, A.C., Cartwright, R.A.: A composite genome approach to identify phylogenetically informative data from next-generation sequencing. BMC Bioinformatics 16(1), 1–10 (2015). 10.1186/S12859-015-0632-Y/FIGURES/7

[23] Tamura, K.: Estimation of the number of nucleotide substitutions when there are strong transition-transversion and G+C-content biases. Molecular Biology and Evolution 9(4), 678–687 (1992). 10.1093/OXFORDJOURNALS.MOLBEV.A040752

[24] Hudson, R.R.: Properties of a neutral allele model with intragenic recombination. Theoretical Population Biology 23(2), 183–201 (1983). 10.1016/0040-5809(83)90013-8

[25] Tavare, S.: Some probabilistic and statistical problems in the analysis of DNA sequences. Lect Math Life Sci (Am Math Soc) 17, 57–86 (1986)

[26] Kozlov, A.M., Darriba, D., Flouri, T., Morel, B., Stamatakis, A.: RAxML-NG: a fast, scalable and user-friendly tool for maximum likelihood phylogenetic inference. Bioinformatics 35(21), 4453–4455 (2019). 10.1093/bioinformatics/btz305

[27] Roch, S., Steel, M.: Likelihood-based tree reconstruction on a concatenation of aligned sequence data sets can be statistically inconsistent. Theoretical population biology 100C, 56–62 (2015). 10.1016/J.TPB.2014.12.005

[28] Chifman, J., Kubatko, L.: Quartet Inference from SNP Data Under the Coalescent Model. Bioinformatics 30(23), 3317–3324 (2014). 10.1093/BIOINFORMATICS/BTU530

[29] Huang, W., Li, L., Myers, J.R., Marth, G.T.: ART: a next-generation sequencing read simulator. Bioinformatics 28(4), 593–594 (2012). 10.1093/BIOINFORMATICS/BTR708

[30] Lefort, V., Desper, R., Gascuel, O.: FastME 2.0: A Comprehensive, Accurate, and Fast Distance-Based Phylogeny Inference Program. Molecular Biology and Evolution 32(10), 2798–2800 (2015). 10.1093/MOLBEV/MSV150

[31] Baumdicker, F., Bisschop, G., Goldstein, D., Gower, G., Ragsdale, A.P., Tsambos, G., Zhu, S., Eldon, B., Ellerman, E.C., Galloway, J.G., Gladstein, A.L., Gorjanc, G., Guo, B., Jeffery, B., Kretzschumar, W.W., Lohse, K., Matschiner, M., Nelson, D., Pope, N.S., Quinto-Cortés, C.D., Rodrigues, M.F., Saunack, K., Sellinger, T., Thornton, K., van Kemenade, H., Wohns, A.W., Wong, Y., Gravel, S., Kern, A.D., Koskela, J., Ralph, P.L., Kelleher, J.: Efficient ancestry and mutation simulation with msprime 1.0. Genetics 220(3), 229 (2022). 10.1093/genetics/iyab229

[32] Sackton, T.B., Grayson, P., Cloutier, A., Hu, Z., Liu, J.S., Wheeler, N.E., Gardner, P.P., Clarke, J.A., Baker, A.J., Clamp, M., Edwards, S.V.: Convergent regulatory evolution and loss of flight in paleognathous birds. Science 364(6435), 74–78 (2019). 10.1126/SCIENCE.AAT7244

[33] Mallo, D., De Oliveira Martins, L., Posada, D.: SimPhy : Phylogenomic Simulation of Gene, Locus, and Species Trees. Systematic Biology 65(2), 334–344 (2016). 10.1093/SYSBIO/SYV082

[34] Fletcher, W., Yang, Z.: INDELible: A Flexible Simulator of Biological Sequence Evolution. Molecular Biology and Evolution 26(8), 1879–1888 (2009). 10.1093/MOLBEV/MSP098

[35] Ge, D., Feijó, A., Wen, Z., Abramov, A.V., Lu, L., Cheng, J., Pan, S., Ye, S., Xia, L., Jiang, X., Vogler, A.P., Yang, Q.: Demographic History and Genomic Response to Environmental Changes in a Rapid Radiation of Wild Rats. Molecular Biology and Evolution 38(5), 1905–1923 (2021). 10.1093/MOLBEV/MSAA334

[36] Ronquist, F., Teslenko, M., Van Der Mark, P., Ayres, D.L., Darling, A., Höhna, S., Larget, B., Liu, L., Suchard, M.A., Huelsenbeck, J.P.: MrBayes 3.2: Efficient Bayesian Phylogenetic Inference and Model Choice Across a Large Model Space. Systematic Biology 61(3), 539–542 (2012). 10.1093/SYSBIO/SYS029

[37] Zhang, C., Rabiee, M., Sayyari, E., Mirarab, S.: ASTRAL-III: Polynomial time species tree reconstruction from partially resolved gene trees. BMC Bioinformatics 19 (2018). 10.1186/s12859-018-2129-y

[38] Wang, S., Teng, D., Li, X., Yang, P., Da, W., Zhang, Y., Zhang, Y., Liu, G., Zhang, X., Wan, W., Dong, Z., Wang, D., Huang, S., Jiang, Z., Wang, Q., Lohman, D.J., Wu, Y., Zhang, L., Jia, F., Westerman, E., Zhang, L., Wang, W., Zhang, W.: The evolution and diversification of oakleaf butterflies. Cell 185(17), 3138–3152 (2022). 10.1016/J.CELL.2022.06.042

[39] Wang, L., DIng, J., Borrell, J.S., Cheek, M., McAllister, H.A., Wang, F., Liu, L., Zhang, H., Zhang, Q., Wang, Y., Wang, N.: Molecular and morphological analyses clarify species delimitation in section Costatae and reveal Betula buggsii sp. nov. (sect. Costatae, Betulaceae) in China. Annals of Botany 129(4), 415–428 (2022). 10.1093/AOB/MCAC001

[40] Wang, N., Kelly, L.J., McAllister, H.A., Zohren, J., Buggs, R.J.A.: Resolving phylogeny and polyploid parentage using genus-wide genome-wide sequence data from birch trees. Molecular Phylogenetics and Evolution 160, 107126 (2021). 10.1016/j.ympev.2021.107126

[41] Patterson, N., Moorjani, P., Luo, Y., Mallick, S., Rohland, N., Zhan, Y., Genschoreck, T., Webster, T., Reich, D.: Ancient admixture in human history. Genetics 192(3), 1065–1093 (2012). 10.1534/GENETICS.112.145037/-/DC1

[42] Corbett-Detig, R.B., Russell, S.L., Nielsen, R., Losos, J.: Phenotypic Convergence Is Not Mirrored at the Protein Level in a Lizard Adaptive Radiation. Molecular Biology and Evolution 37(6), 1604–1614 (2020). 10.1093/MOLBEV/MSAA028

[43] Kozlov, A.M., Aberer, A.J., Stamatakis, A.: ExaML version 3: a tool for phylogenomic analyses on supercomputers. Bioinformatics 31(15), 2577–2579 (2015). 10.1093/BIOINFORMATICS/BTV184

[44] He, S., Li, L., Lv, L.Y., Cai, W.J., Dou, Y.Q., Li, J., Tang, S.L., Chen, X., Zhang, Z., Xu, J., Zhang, Y.P., Yin, Z., Wuertz, S., Tao, Y.X., Kuhl, H., Liang, X.F.: Mandarin fish (Sinipercidae) genomes provide insights into innate predatory feeding. Communications Biology 2020 3:1 3(1), 1–13 (2020). 10.1038/s42003-020-1094-y

[45] Stamatakis, A.: RAxML version 8: a tool for phylogenetic analysis and post-analysis of large phylogenies. Bioinformatics 30(9), 1312–1313 (2014). 10.1093/BIOINFORMATICS/BTU033

[46] Wang, G.-S., Cai, Q., Hao, Y.-J., Bau, T., Chen, Z.-H., Li, M.-X., David, N., Kraisitudomsook, N., Yang, Z.-L.: Phylogenetic and taxonomic updates of Agaricales, with an emphasis on Tricholomopsis. Mycology, 1–30 (2023). 10.1080/21501203.2023.2263031

[47] Shao, Y., Zhou, L., Li, F., Zhao, L., Zhang, B.L., Shao, F., Chen, J.W., Chen, C.Y., Bi, X., Zhuang, X.L., Zhu, H.L., Hu, J., Sun, Z., Li, X., Wang, D., Rivas-González, I., Wang, S., Wang, Y.M., Chen, W., Li, G., Lu, H.M., Liu, Y., Kuderna, L.F.K., Farh, K.K.H., Fan, P.F., Yu, L., Li, M., Liu, Z.J., Tiley, G.P., Yoder, A.D., Roos, C., Hayakawa, T., Marques-Bonet, T., Rogers, J., Stenson, P.D., Cooper, D.N., Schierup, M.H., Yao, Y.G., Zhang, Y.P., Wang, W., Qi, X.G., Zhang, G., Wu, D.D.: Phylogenomic analyses provide insights into primate evolution. Science 380(6648), 913–924 (2023). 10.1126/SCIENCE.ABN6919/SUPPL{}FILE/SCIENCE.ABN6919{}MDAR{}REPRODUCIBILITY{}C

